# Basement membrane sets the timescale of tissue mechanical memory

**DOI:** 10.64898/2026.04.23.720416

**Authors:** Nargess Khalilgharibi, Billie Meadowcroft, Anđela Šarić, Yanlan Mao

## Abstract

The basement membrane (BM) is a specialised extracellular matrix tightly tethered to epithelial tissues. While the viscoelastic response of epithelial cells to external deformation has been widely studied, the dynamic mechanical role of its underlying BM remains poorly understood. This is mainly due to its thin, dense, non-fibrillar structure and limited number of model systems that allow live fluorescent imaging of the BM components. Using the *Drosophila* wing disc, we investigate the BM’s response to sustained deformation and find that the tissue retains memory of its shape for up to four hours, enabled by the BM’s initial elasticity. However, prolonged deformation leads to BM network rearrangement and loss of mechanical memory, resulting in permanent shape change. Our findings reveal that the BM sets the long-term viscoelastic timescale of epithelial tissues which plays a critical role in maintaining tissue architecture under mechanical stress.

## Main

The basement membrane (BM) is a specialised extracellular matrix that is tightly adhered to the basal side of epithelial tissues. During development and in normal physiology, epithelial tissues are exposed to different magnitudes^1–4^, rates^1,2,5,6^ and durations^7^ of deformations. While in some processes (especially those during development and growth) tissues undergo irreversible deformations, in many other instances they must revert to their original shape after deformations, which we define here as their mechanical memory.

The response of a tissue to deformation depends on the mechanical properties of the cells and their tightly tethered BM. Over short timescales, it has been shown that epithelial tissues act as a viscoelastic solid in response to external deformation^8^. The actomyosin cytoskeleton is a key player in this mechanical behaviour, for example through dictating the rate of stress dissipation^9^ and generating supracellular cables to provide tissue-scale elasticity^10^. At longer timescales, cellular processes such as neighbour exchange^11–13^ and oriented cell divisions^14,15^ help the tissue to dissipate the stress further while resulting in irreversible tissue shape change.

While the mechanical response of epithelial cells to deformation has been studied widely, little is known about the BM response to deformation^16,17^. Indeed, studies characterising the mechanical properties of extracellular matrices have mainly focused on the interstitial matrix of soft tissues^18^. Unlike interstitial matrices which are predominantly composed of fibrillar collagens, BM is a thin dense non-fibrillar structure^19^, making its characterisation and mechanical manipulation experimentally challenging. Recent work has shed light on the static mechanical properties of BM *ex vivo*^20–23^. However, the characterisation of the dynamic mechanical properties is still limited to isolated BMs^24^ and reconstituted matrices derived from animal BMs (e.g. Matrigel)^25,26^.

Here, we study the dynamic response of BM to deformation and its role in maintaining tissue shape. We expose the *Drosophila* wing disc epithelia to several hours of sustained external deformation. We find that the tissue can maintain memory of its shape for up to 4 hours, much longer than the timescales reported for cellular aggregates^27,28^ and epithelial monolayers devoid of a substrate^8,29^. We find that the BM provides the tissue with this long memory of shape. Our analysis shows that the BM deforms elastically in response to deformation. Remarkably, this elasticity is sustained for several hours, which gives the tissue a prolonged mechanical memory, allowing it to revert to its original shape when stress is removed, up to 4 hours later. If the deformation is maintained, the BM network gradually re-arranges over the course of 4 hours, leading to gradual loss of memory of its initial shape as it adapts to the new deformed shape. Finally, we adopt a coarse-grained model of collagen IV network, one of the main components of the BM, to link the macroscopic changes observed experimentally to the molecular changes in the network. Together, we show that the BM sets the long-term viscoelastic timescale of the tissue, providing it with mechanical robustness against external deformations.

## Results

### The BM helps tissues maintain memory of their shape over long periods of stretch

To characterise how long tissues can maintain memory of their shape, we exposed the *Drosophila* wing disc epithelium to different durations of stretch (Fig. 1a). To measure the stress sustained in the tissue and assess whether the tissue had relaxed the stress as it adapts to the stretched shape, we performed apical annular laser ablation cuts^30^ and quantified the aspect ratio of the outer boundary of the cut after different durations of stretch (Fig. 1a-f, Fig. S1a-e). Upon annular ablation, the inner and outer boundaries of the annular cut recoil in opposite directions. If the tissue is under isotropic tension, the boundaries will recoil with the same rate in all directions, leading to a final circular shape of both inner and outer boundaries (Fig. 1b). However, if the tissue is under anisotropic tension, the tissue will recoil more in the direction of higher tension, leading to a final elliptical shape of inner and outer boundaries^30^ (Fig. 1b-c). We quantify the outer ellipse aspect ratios (Fig. 1c). When we performed annular cuts on wing discs mounted in the stretcher without being stretched (anchor, Fig. 1d-f), we observed outer ellipse aspect ratios of 1.08 ± 0.04, suggesting a relatively isotropic tension when the disc is simply anchored in the stretcher. Upon stretch, the aspect ratio of the outer ellipse, performed within 30min of stretch, increased to 1.25 ± 0.09, suggesting significantly higher anisotropic tension than the anchor (Fig. 1e,f). When we performed annular cuts on discs at successively longer periods of stretch (Fig. 1f), we found the tissue was able to maintain the higher aspect ratio for the first 4h of stretch. After 5h of maintained stretch, the aspect ratios of the cuts returned to values similar to the anchor (1.11 ± 0.04) (Fig. 1e,f), suggesting that the stress was relaxed. To ensure that this was due to the adaption of the tissue to the stretched shape, and not due to the viability of the wing discs in the stretcher, we mounted the wing discs in the stretcher for 5h, before applying a stretch and cutting immediately after. The aspect ratios of the cuts were similar to those of wing discs stretched and cut immediately after being mounted, suggesting that the 5h mounting in the stretcher was not affecting the viability or response of the tissue (Fig. S1h).

**Fig 1:**
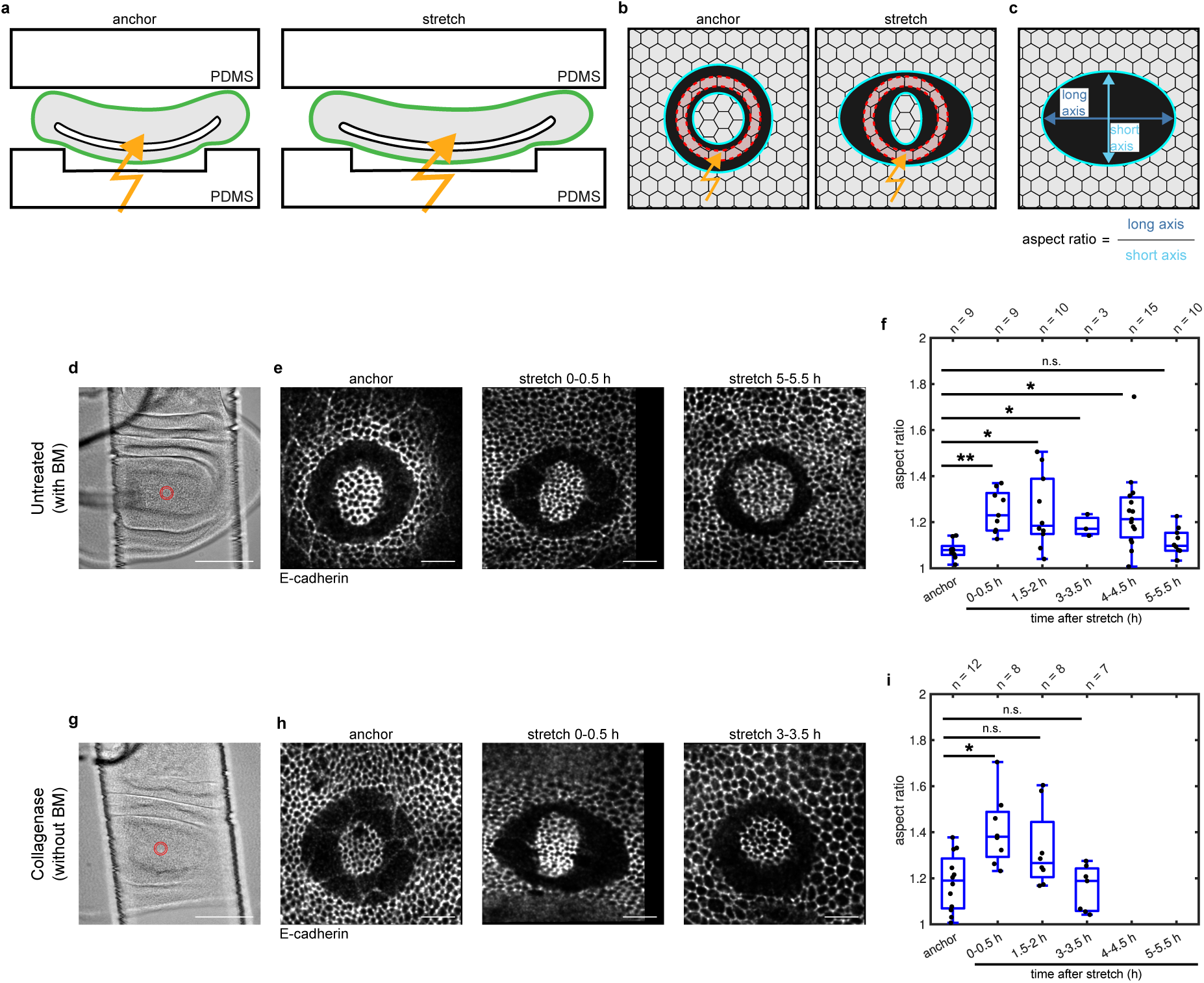
The BM helps the tissue to maintain memory of its shape over long periods of stretch. (a) Schematic diagram of the annular ablation experiments in our custom-made stretcher. A wing disc (consisting of epithelium in grey and BM in green) is sandwiched between two layers of PDMS that are clamped on our custom-made stretcher. Annular ablations are performed with a multiphoton laser shining from the bottom (orange arrow). We therefore ensure that the wing discs are mounted with the apical side of the columnar layer facing down, therefore being closer to the laser. Ablations are done in “anchor” (i.e. when the wing disc is mounted on the stretcher but not stretched, left panel) and “stretch” (right panel) conditions. (b) Schematic diagram of an annular ablation and quantification of aspect ratio. An annular region (red) is ablated with the laser (orange arrow). The inner and outer boundaries of the annular region recoil inwards and outwards respectively (cyan). In this paper, we quantify the aspect ratio of the outer boundary. In absence of external stretch (anchor, left panel) the recoil is homogenous in all directions, resulting in a circle-like boundary. Upon stretch, the recoil is larger in the direction of stretch, resulting in an ellipse-like boundary (right panel). (c) The outer boundary of the annular cut is fitted with an ellipse and the aspect ratio is calculated. (d) Example brightfield image of an untreated stretched wing disc in the PDMS channel. The ablated region is also shown (red circles). Scale bar = 100 μm. (e) Snapshot of the annular ablation experiments of untreated wing discs in anchor (left panel) and stretched (0-0.5 h: middle panel; 5-5.5 h: right panel) conditions. The cell boundaries are marked with DE-cad-GFP. Scale bars = 10 μm. (f) Box plots comparing aspect ratio of annular ablations of wing discs subject to different durations of stretch with anchored wing discs (** P < 0.001, Statistical power > 80% at α = 0.01 for 0-0.5 h stretch; * P = 0.003, Statistical power > 80% at α = 0.05 for 1.5-2 h stretch; * P = 0.02, Statistical power > 80% at α = 0.05 for 3-3.5 h stretch; * P = 0.004, Statistical power > 80% at α = 0.05 for 4-4.5 h stretch; n.s., P = 0.32 for 5-5.5 h stretch; all compared with anchor). (g) Example brightfield image of collagenase-treated stretched wing disc in the PDMS channel. The ablated region is also shown (red circles). Scale bar = 100 μm. (h) Snapshot of the annular ablation experiments of collagenase-treated wing discs in anchor (left panel) and stretched (0-0.5 h: middle panel; 5-5.5 h: right panel) conditions. The cell boundaries are marked with DE-cad-GFP. Scale bars = 10 μm. (i) Box plots comparing aspect ratio of annular ablations of anchored wing discs with those subject to different durations of stretch (* P = 0.005, Statistical power > 80% at α = 0.05 for 0-0.5 h stretch; n.s., P = 0.11 for 1.5-2 h stretch; n.s., P = 0.77 for 3-3.5 h stretch; all compared with anchor). All wing discs were treated with collagenase prior to the experiment.

It has previously been shown that epithelial tissues devoid of a substrate adapt to the stretched shape in timescales of less than 1 minute^9^. We therefore asked whether the 5h timescale that we observed in the wing discs is due to the presence of the BM. To test this, we removed the BM through enzymatic digestion using collagenase (Fig. S1f-g). When we stretched the collagenase-treated wing discs, and performed annular cuts shortly after stretch, we found that the outer ellipse aspect ratio increased (1.41 ± 0.15 compared to 1.18 ± 0.12 for anchor), like untreated wing discs (Fig. 1g-i). However, this aspect ratio reduced and was no longer significantly different to anchored wing discs when the cut was performed after 1.5h of stretch, and completely disappeared after 3h of stretch (Fig. 1i), suggesting faster stress relaxation when discs do not have a BM. This suggests that the BM helps the tissue to maintain a longer mechanical memory of its shape. In other words, the tissue loses its elasticity faster without the BM.

### The BM relaxes the applied stress over time by changing its network organisation

To understand how the BM helps the tissue to maintain elasticity, we asked how does the BM itself respond to stretch? To answer this, we repeated previous experiments of exposing the tissue to different durations of stretch, but this time we performed the annular cuts on the BM only, taking care not to damage the cells (Fig. 2a-c, Fig. S2a). The aspect ratio of the outer boundary of the cut, in the anchor case, was similar to the aspect ratio on the apical side (1.12 ± 0.10, Fig. 2d). Upon short stretch, the aspect ratios of the BM cut increased to 1.19 ± 0.09, again similar to that on the apical surface. This suggests that the tissue tension previously measured apically is also borne by the BM. We further observed that if the stretch is maintained for 3h, the aspect ratios of the BM annular cuts remained elevated and were only significantly reduced after 4h (Fig. 2e), suggesting that the BM requires 4h to relax the elevated stretch-induced tension. This is considerably longer than the time scale of stress relaxation when the BM is removed (Fig. 1i)

**Fig 2:**
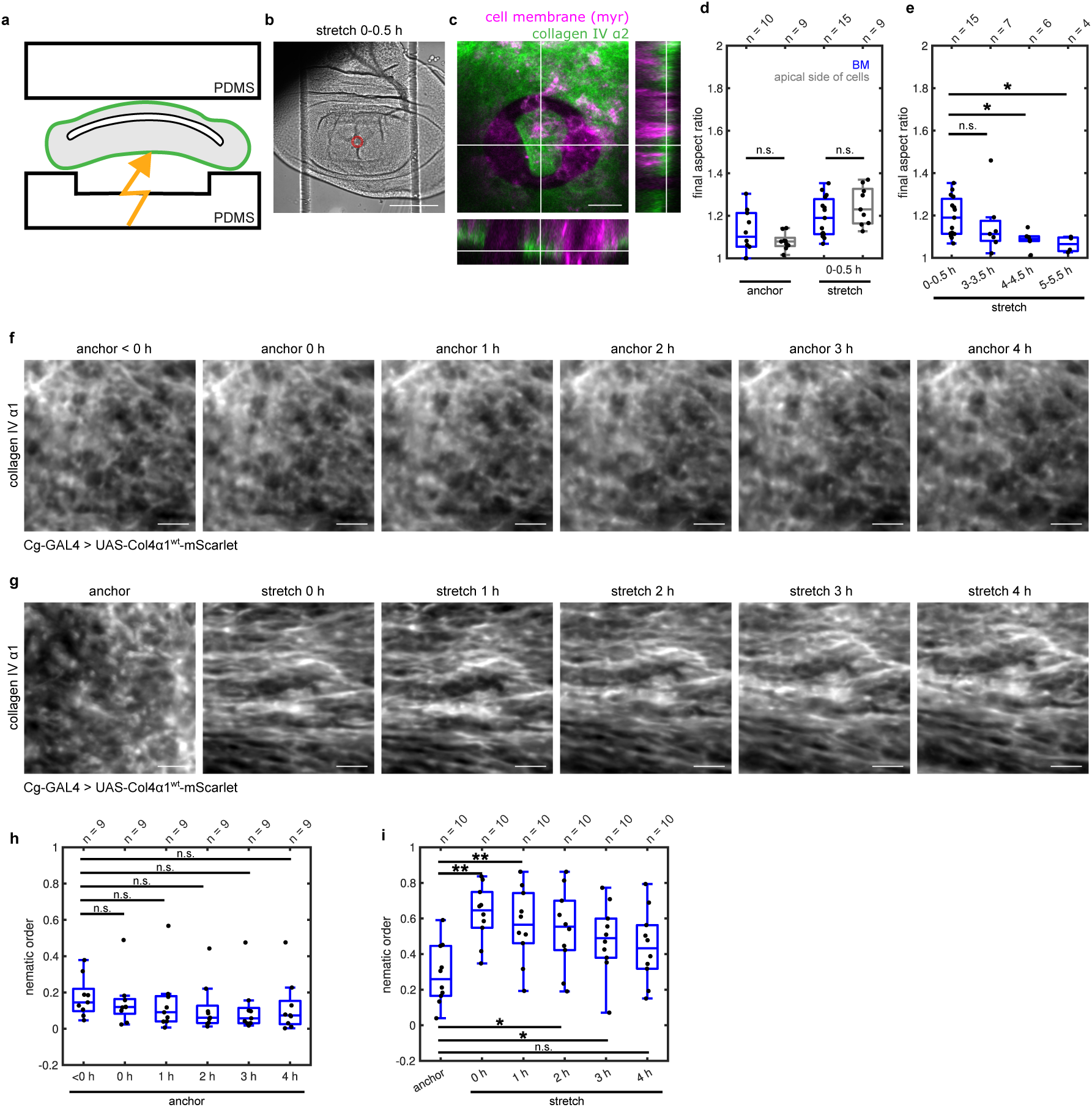
The organisation of the collagen IV network changes over time in response to sustained external deformation. (a) Schematic diagram of the annular ablation experiments of the BM (epithelium: grey, BM: green, laser: orange). Wing discs are mounted such that the BM of the columnar layer is facing down, therefore being closer to the laser. (b) Example brightfield image of an untreated stretched wing disc with in the PDMS channel. The ablated region is also shown (red circles). Scale bar = 100 μm. (c) Zoomed-in confocal microscopy image of the ablated region in (b) with cell membranes (*myr*) in magenta and BM (Col4α2) in green. Scale bar = 10 μm (right). (d) Comparison of aspect ratios of annular cuts at the BM and apical surface of the columnar layer (n.s., P = 0.50 for anchor; n.s., P = 0.15 for 0-0.5 h stretch). (e) The aspect ratios of the outer boundary of the annular cuts of the BM in wing discs subjected to different durations of stretch (n.s., P = 0.23 for 3-3.5 h stretch; * P = 0.01, Statistical power > 80% at α = 0.05 for 4-4.5 h stretch; * P = 0.004, Statistical power > 80% at α = 0.05 for 5-5.5 h stretch; all compared to 0-0.5 h stretch). (f,g) Near-super-resolution confocal images of the collagen IV network (Col4α1) in anchored (f) and stretched (g) conditions. The field of view shows a 250 × 250 pixel (27.08 μm × 27.08 μm) region that is used for the analysis of the nematic order. Scale bars = 5 μm. (h,i) Box plots of the change in nematic order over time for anchored (h) and stretched (i) wing discs. (In panel h: n.s., P = 0.16 for 0 h anchor; n.s., P = 0.16 for 1 h anchor; n.s., P = 0.10 for 2 h anchor; n.s., P = 0.13 for 3 h anchor; n.s., P = 0.16 for 4 h anchor; all compared with <0’ anchor. In panel i: ** P = 0.002, Statistical power > 80% at α = 0.01 for 0 h stretch; ** P = 0.002, Statistical power > 80% at α = 0.01 for 1 h stretch; * P = 0.004, Statistical power > 80% at α = 0.05 for 2 h stretch; * P = 0.010, Statistical power > 80% at α = 0.05 for 3 h stretch; n.s., P = 0.02, Statistical power = 60% at α = 0.05 for 4 h stretch; all compared with anchor).

How does the BM relax the tension? For this, we focused on the structure of collagen IV network, which is one of the main components of the BM^19^. Polymer networks can relax stress through various mechanisms including breaking and forming bonds^31^. This can involve rearrangement of existing bonds (without incorporation of new material), or the removal of existing molecules and incorporation of new molecules into the network. In biological systems, the latter is one of the main contributors to stress relaxation^29,32–34^. In the wing disc, collagen IV is produced externally in the Fat Body and then transported through the haemolymph to the wing disc^35^. Therefore, when the wing discs are dissected and cultured *ex vivo,* as we do here, there is no source of new collagen IV and previous Fluorescence Recovery After Photobleaching (FRAP) experiments have shown that there is no addition of new material in the network^22,23^ (Fig. S2c).

To understand whether the rearrangement of existing bonds could be the source of stress relaxation in the BM, we asked how the collagen IV network organisation changes over time. For this, we stretched wing discs overexpressing mScarlet-tagged collagen IV under the control of the collagen IV promoter and imaged them over the course of 4h using near super-resolution live microscopy (Fig. 2f-g, Fig. S2d). We measured the global alignment in the network, quantified by the nematic order, using the Alignment by Fourier Transform (AFT) workflow^36^. We found that immediately upon stretch, the network becomes more aligned (Fig. 2h-i). This observation supports the common view in the field that BM network behaves elastically, at least over short timescales. However, as the external deformation is maintained, the alignment gradually goes down, with values becoming not significantly different from anchor after 4h of stretch (Fig. 2h-i). Since the timescale of this rearrangement without inclusion of new material is comparable to the timescales of tension relaxation in the BM (Fig. 2e), this suggests that the re-arrangement of the existing BM network results in the relaxation of the tension.

### A coarse-grained model of the collagen IV network

The AFT analysis gave us an understanding of the temporal changes in the collagen IV network at the macroscopic level. However, even though the images are at the limits of current live microscopy technology, they do not provide molecular-level understanding of how the network is re-arranging to relax tension. We therefore developed a coarse-grained molecular dynamics simulation of the collagen IV network to understand the physical mechanism behind the macroscopic changes we observe. The model is based on our previously published model^37^. Briefly, we modelled the collagen IV protomer as a two-bead rod, with one bead representing the NC1 domain and the other bead representing the 7S domain of the collagen IV molecule. In accordance with the literature^19^, the NC1 bead can bind to only one other NC1 bead, while the 7S bead can bind to up to three other 7S beads (Fig. 3a). These protomers self-assemble to form a percolating network (Fig. 3b). Since in our experiments, the mScarlet fluorescent protein is inserted close to the 7S domain^38^, we hypothesised that experimentally we are visualising only one end of the collagen IV molecule. Therefore, in our simulations, we also only “visualise” the 7S beads. Furthermore, since the fluorescent collagen IV is overexpressed in our experiments in addition to the endogenous non-fluorescent collagen IV, we only visualised a sub-population of the protomers in our simulations (methods). Finally, considering the resolution limit of the microscope, we applied a Gaussian spread function to the 7S beads, so that we could compare the nematic order of simulation snapshots with those of microscopy images using the AFT workflow (Fig. 3c).

**Fig 3:**
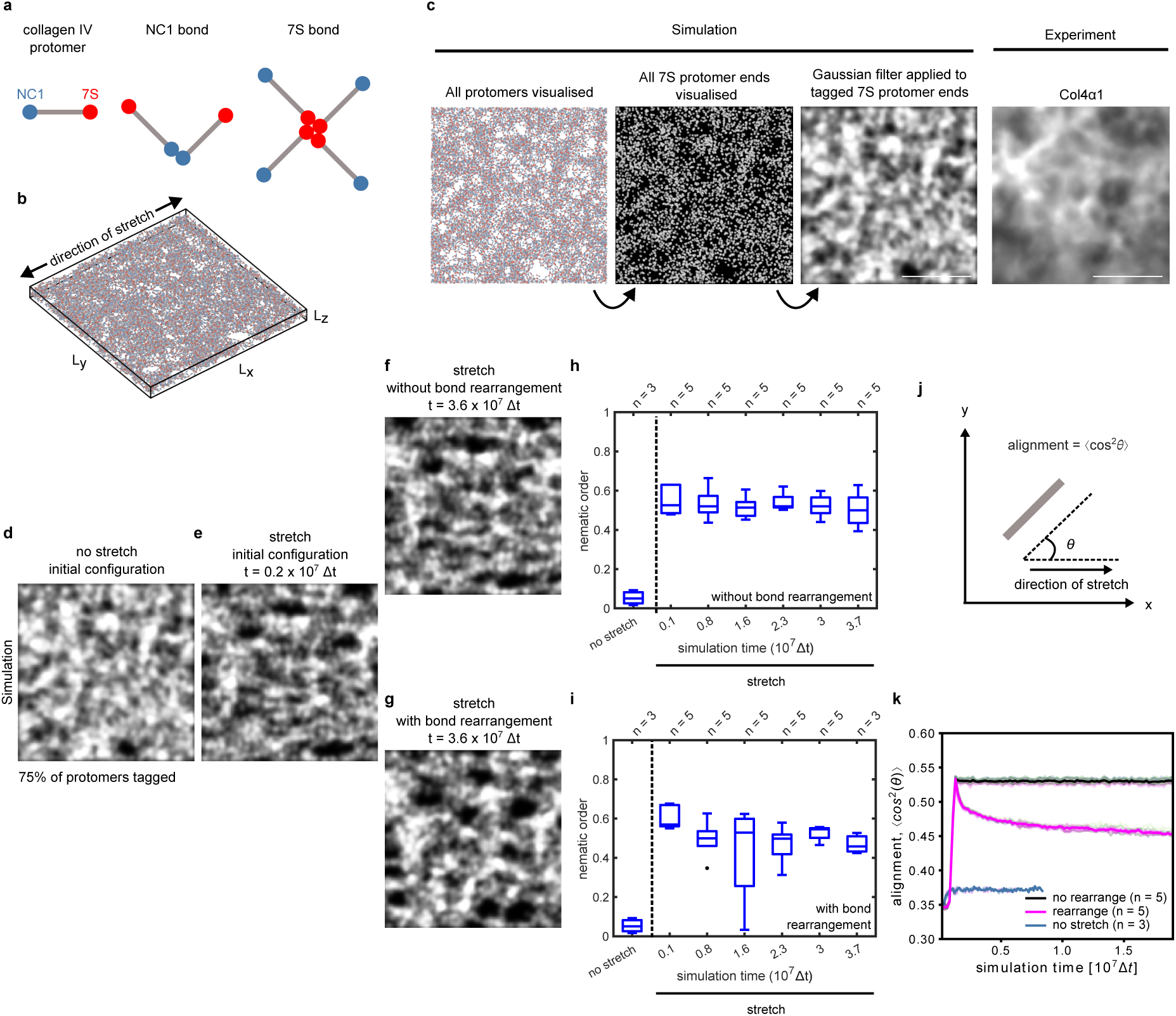
A coarse-grained model of collagen IV network captures temporal changes in its organisation upon stretch. (a) The collagen IV protomer is modelled as a two-bead rod (left panel). The protomer can form dimers at the NC1 domain (blue beads, middle panel) and tetramers at the 7S domain (red beads, right panel). (b) The protomers self-organise into a network. The dimensions of the simulation box are represented as ***L*_*x*_ = *L*_*y*_ = 164σ** and ***L*_*z*_ = 12σ**. The simulation length unit σ corresponds to 80 nm. (c) Steps to transform simulation snapshots to those comparable with microscopy images. The left panel shows a raw snapshot of a simulation with all protomers visualised. In the 2^nd^ left panel, only the 7S protomer ends are shown. In the 2^nd^ right panel, a Gaussian filter is applied to the tagged 7S protomer ends. The right panel shows a zoomed-in region from Fig. 2f anchor. Scale bars = 5 μm. (d-g) Snapshots of two simulations with normal protomers and without/with remodelling over time. Only 75% of the protomers are tagged to replicate experimental conditions. The initial configurations before and immediately after stretch are the same for both simulations (d,e). Then, one simulation continues without bond rearrangement (f) while bond rearrangement is enabled for the other simulation (g). (h,i) Box plots showing temporal evolution of the nematic order of simulated networks without (h) and with (i) bond rearrangement. (j) Schematic diagram of the protomer alignment. (k) Temporal evolution of the protomer alignment in different simulation conditions (without remodelling: black, with remodelling: magenta, no stretch: blue). Individual simulations are shown half-transparent with different colours. The solid lines show average of multiple simulations.

After obtaining the self-assembled network, we allowed it to equilibrate (Fig. S3a, cream region) before applying a stretch. This represents our simulated “no stretch initial configuration” (Fig. 3d). Next, we stretched the simulated network uniaxially and assessed its response (Fig. S3f,g). To calibrate the extent of deformation in simulations, we simulated annular ablations on stretched simulated networks and hereafter chose an applied deformation which matched the anisotropic recoil response recorded in the experiments (Fig. S3c, methods). In the model we could implement bond rearrangement, i.e. stochastic bond breaking and formation without introducing any new protomers into the network, simulating our experimental *ex-vivo* condition where there is no addition of new material. We first asked how the network would respond in absence of bond rearrangement. We found that upon stretch, the nematic order did increase, but unlike experiments, it remained elevated (Fig. 3e,f,h). When we analysed the amount of stress in the network (Fig. S3h), we observed a rapid relaxation that plateaued at ∼ 83% of the maximum stress. By fitting an exponential function to this relaxation, we estimated its characteristic timescale as τ_1_ ∼ 6 × 10^4^ Δ*t*, where Δ*t* is the simulation timestep (Fig. S3a, dark green region). To relate the simulation and experimental timescales, we compared the plateauing of annular ablations in both contexts (Fig. S3b, methods). We estimated the simulation timescale of τ_2_∼ 1.5 × 10^6^ Δ*t* (Fig S3b) to be equivalent to the minute timescale of experimental ablations (Fig. S2b). Since τ_2_ ≫ τ_1_ And τ_2_ is equivalent to a minute in our experiments, we conclude that we cannot observe this rapid relaxation experimentally, because all our measurements are taken several minutes after stretch. Together, these findings show that upon stretch, the network can relax the stress without breaking bonds and solely through repositioning of existing bonds. However, this relaxation, likely driven by thermal fluctuations, is very fast (i.e. sub-minute) and therefore cannot account for the slow, multi-hour changes in nematic order observed experimentally.

Next, we introduced bond rearrangement, where all bonds are allowed to break with a constant probability if they stretch beyond a threshold length. With bond rearrangement, we observed a trend of temporal reduction in nematic order, similar to the experiments (Fig. 3g,i). Since in the experiments, only the 7S domain of the collagen IV molecule is tagged, we cannot observe the whole molecule and therefore the whole meshwork. What seems like a connected meshwork in the microscopy images only appears as such due the fluorescent limit of the microscope, as well as overlaying of multiple layers of BM^39^. We therefore took advantage of the simulations to directly quantify the alignment of the collagen IV protomers relative to the axis of stretch (Fig. 3j). Upon stretch, the molecular alignment increases as expected, however, it only decreases in presence of bond rearrangement (Fig. 3k). A similar pattern was observed for the network stress (Fig. S3h). Together, these simulations show that the change in the collagen IV nematic order we observe experimentally is likely a result of bond rearrangement, which relaxes stress in the network.

### A well-connected BM network is required for its effective response to stretch

Bond rearrangement results in changes to network connectivity. Having shown that the collagen IV network can relax external stress through bond rearrangement, we then asked what would happen if we perturbed the connectivity of the network. To do this, we modified the simulations such that a portion of the 7S beads – the same population that were tagged and visualised in simulations – could not form bonds. We run two sets of simulations, one with 75% and one with 56% mutant protomers (methods). In both scenarios, we found that the perturbed network did not respond as much as the non-perturbed network to stretch, with the network nematic order, molecular alignment and stress only marginally increasing upon stretch (Fig. 4a-d, S3i-j). While the nematic order remained largely unchanged after stretch (Fig. 4c), both molecular alignment and stress decreased (Fig. 4d, S3j compared to Fig. 3k, S3h). To test the model prediction experimentally, we used wing discs overexpressing an mScarlet-tagged collagen IV α1 mutant under the control of the collagen IV promoter (Fig. 4e-hThis lacks part of the putative 7S domain and has been shown to affect collagen IV network assembly^38^. We found that the network did not respond effectively to stretch, with the nematic order not changing significantly upon and throughout the 4 h stretch (Fig. 4e-h). This agrees with our simulations, suggesting that a well-connected network is required for an effective response to stretch.

**Fig 4:**
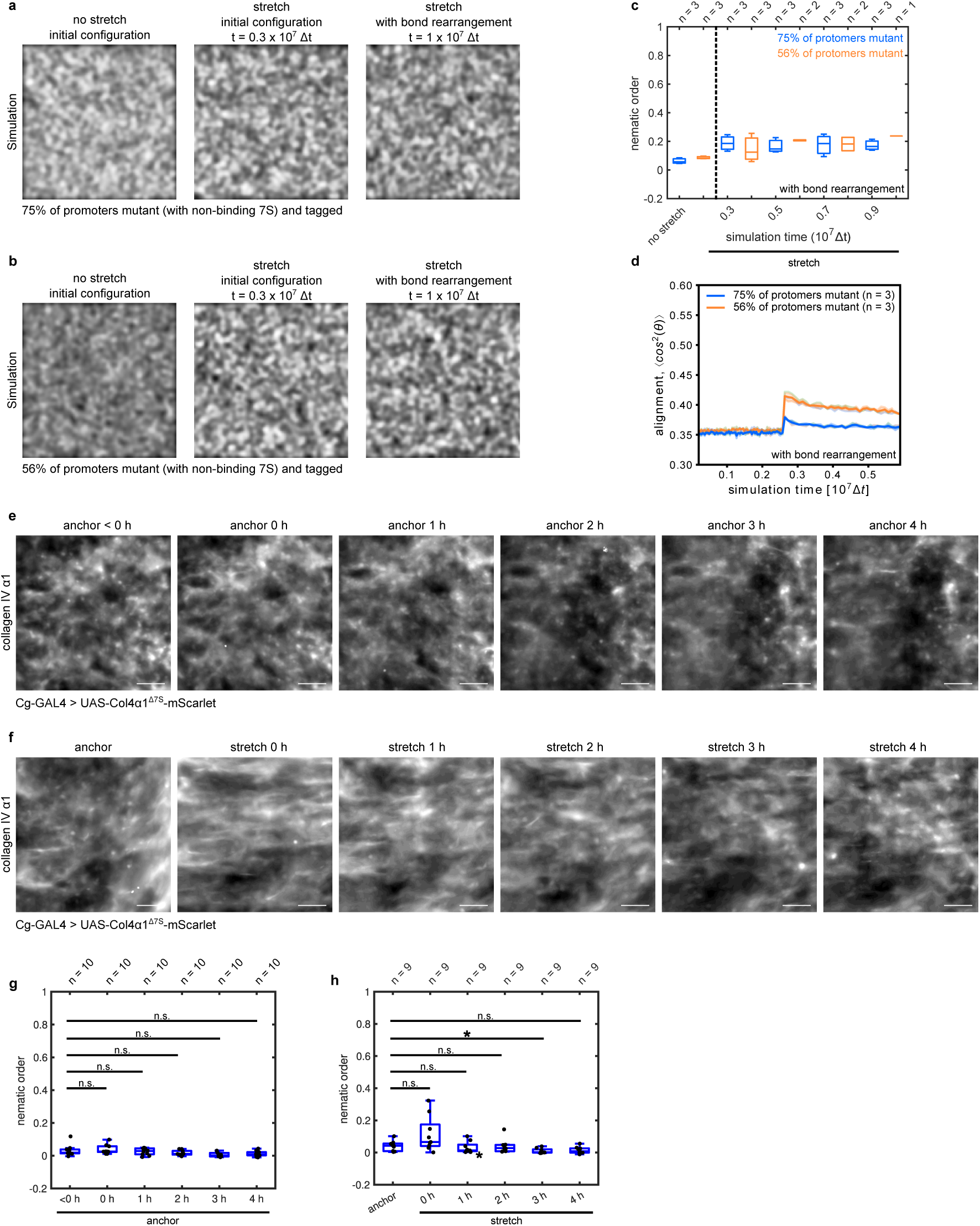
Having a sub-population of mutant collagen IV molecules perturbs the mechanical response of the network to deformation. (a,b) Snapshots of stretching simulations with mutant protomers and bond rearrangement over time. Two different concentrations of mutant protomers were tested. Timepoints before (no stretch, left panel), shortly after (middle panel) and a long time after (right panel) stretch are shown. (c) Box plot showing temporal evolution of the nematic order of simulated networks with bond rearrangement for two different concentrations of mutant protomers. (d) Temporal evolution of the protomer alignment upon stretch for simulations with different concentrations of mutant protomers. Individual simulations are shown half-transparent with different colours. The solid lines show average of multiple simulations. (e,f) Near-super-resolution confocal images of the collagen IV network (Col4α1) of mutant wing discs in anchored (e) and stretched (f) conditions. Scale bar = 5 μm. (g,h) Box plots of the change in nematic order over time for anchored (f) and stretched (g) mutant wing discs. (In panel g: n.s., P = 0.70 for 0 h anchor; n.s., P = 0.92 for 1 h anchor; n.s., P = 0.38 for 2 h anchor; n.s., P = 0.004, Statistical power = 78% at α = 0.05 for 3 h anchor; n.s., P = 0.32 for 4 h anchor; all compared with <0 h anchor. In panel h: n.s., P = 0.02, Statistical power = 65% at α = 0.05 for 0 h stretch; n.s., P = 0.57 for 1 h stretch; n.s., P = 0.73 for 2 h stretch; * P = 0.01, Statistical power > 80% at α = 0.05 for 3 h stretch; n.s., P = 0.05 for 4 h stretch; all compared with anchor).

## Discussion

The mechanical response to deformation of complex multi-component materials often has various timescales that can be attributed to its components^7,40^. Here, we characterise the dynamic mechanical response of BM, the specialised ECM layer lining epithelial tissues, to deformation and show that it dictates the tissue stress relaxation at multi-hour timescales. This longer relaxation timescale of BM compared to cells allows it to act as a “mechanical memory” for the tissue, helping it to maintain its shape in response to transient deformations.

Breaking and reforming of bonds leads to stress relaxation in polymer networks^31^. Biologically, this is often associated with the incorporation of new polymer subunits and the degradation of existing ones. However, it is possible to relax the stress through rearrangement of existing bonds^25,41^. Here, we took advantage of the *Drosophila* wing disc system, where we can visualise the BM network and where there is no addition of new material *ex viv*o. This allowed us to explore stress relaxation through rearrangement of existing network bonds, which is less well-characterised in biological contexts. The current resolution limits of light microscopy do not allow imaging of the BM dynamics at molecular resolution. We therefore adapted our previously published molecular dynamics model^37^ to create simulation images comparable to experiments. This helps to bridge the gap between microscopic resolution of confocal imaging and sub-micron network structures, by allowing validation of simulation results directly with measurable experimental quantities (e.g. nematic order), while also adding more insight into the molecular mechanisms behind it.

Collagen IV forms a highly connected network, with the two main NC1 and 7S bonds at the molecule ends, as well as non-covalent lateral associations along the length of the of the molecules and further crosslinking through other BM components. Here, we used mutations of the 7S domain that affect the BM integrity to demonstrate the importance of having a well-connected network in the BM mechanical response. Indeed, the higher the proportion of the mutant protomers in the model, the less stress was taken up by the BM. Recent studies have highlighted how other components of the BM network contribute to its mechanical properties, for example through affecting its structural integrity^44,45^ and turnover rate^34^. The model can be adapted to incorporate these additional components. Together, we have developed and used a pipeline combining near-super-resolution imaging with molecular dynamics modelling to better understand the dynamic changes in the BM structure that contributes to the viscoelastic mechanical response of tissues to stress. Using this pipeline in model organisms such as *Drosophila* or *C-elegans* where a large library of tagged BM proteins exists^46^ will provide a versatile framework to characterise the dynamic mechanical properties of individual BM components without the need to use animals or widely-used animal-derived BMs (e.g. Matrigel) and ultimately help engineer smart synthetic matrices that better capture mechanical properties of natural BMs.

## Methods

### *Drosophila* melanogaster strains

*Drosophila* stocks were raised on conventional cornmeal media at 25°C. Male and female larvae were dissected at mid-late 3^rd^ instar development (96-120 h AEL) for experiments. *Ex vivo* culture media consisted of Shields and Sang M3 media (Sigma) supplemented with 2% FBS (Sigma), 1% pen/strep (Gibco), 3ng/ml ecdysone (Sigma) and 2ng/ml insulin (Sigma). For stretching experiments, wing discs were mounted in our custom-made stretcher, as described by Duda et al^10^ and briefly explained below under Methods: Stretching experiments. The alleles and transgenes used are shown in Table 1. The experimental genotypes and their corresponding data are shown in Table 2.

**Table 1:** List of alleles and transgenes.

| Allele or transgene | Source | FlyBase ID |
| --- | --- | --- |
| DE-cad-GFP | Huang *et al*<sup>47</sup> | FBti0168565 |
| *pnr*-GAL4 | BDSC 3039 | FBtp0090138 |
| *sqh*-mCherry | Martin et al<sup>48</sup> | FBtp0065864 |
| *vkg*-GFP | BDSC 98343 | FBti0153267 |
| CAAX-RFP | DGRC 109825 | NA |
| *tub*-GAL4 | BDSC 5138 | FBti0012687 |
| *Cg*-GAL4 | Pastor-Pareja & Xu<sup>35</sup> | FBtp0012452 |
| UAS-Col4α1<sup>wt</sup>-mScarlet | Gift from the Stramer Lab<sup>38</sup> | NA |
| UAS-Col4α1<sup>Δ7S</sup>-mScarlet | Gift from the Stramer Lab<sup>38</sup> | NA |
| UAS-*myr*-mRFP | BDSC 7118 | FBti0027895 |
| *Act5C*-GAL4 | BDSC 4414 | FBtp0001406 |

**Table 2:** List of *Drosophila* experimental genotypes and corresponding figures.

| Genotype | Corresponding Figures and Videos |
| --- | --- |
| *w*; DE-cad-GFP; *pnr*-GAL4, *sqh*-mCherry | Fig. 1d-i, S1b-e,h |
| *w*; *vkg*-GFP, CAAX-RFP; *tub*-GAL4/TM6B | Fig. S1f-g |
| *yw/w*; *vkg*-GFP, UAS-*myr*-mRFP/*vkg*-GFP; *Act5C*-GAL4/+ | Fig. 2b-e, S2a-b |
| *w*; *Cg*-GAL4/+; UAS-Col4α1<sup>wt</sup>-mScarlet/+ | Fig. 2f-i, S2d |
| *w*; *vkg*-GFP, UAS-*myr*-mRFP; *Act5C*-GAL4/TM6B | Fig. S2c |
| *w*; *Cg*-Gal4/+; UAS-Col4α1<sup>Δ7S</sup>-mScarlet /+ | Fig. 4e-h |

### Drug treatments

To remove BM, a collagenase solution was prepared by mixing an appropriate amount of collagenase type-I (Worthington) with the culture media to reach a final activity of 624 units per ml. Dissected wing discs were first incubated in 50% concentration of the collagenase solution for 5 minutes, then mounted in the stretcher, and finally incubated in full concentration of collagenase for another 15 minutes. The total incubation time in collagenase was 30 minutes.

### Stretching experiments

A custom-made stretcher was used, as described in Duda et al^10^. Briefly, the stretcher consisted of two metal arms that are connected to and can be moved apart using a fine pitch screw. A pre-patterned polydimethylsiloxane (PDMS) membrane was spread over the two arms, overlaid with an unpatterned PDMS membrane (GelPak, 6.5 mil) and clamped on both ends. Next, 5-10 isolated wing discs suspended in the culture media were injected in between the two PDMS layers and positioned with a micro-dissecting needle over a microchannel, with the appropriate side of wing disc proper facing down the channel (i.e. apical side for imaging the apical side and basal side for imaging the BM). A 500 ml drop of culture media was placed on top of the membranes and topped up every hour during the experiments.

The pre-patterned PDMS membranes were made as follows: PDMS was prepared by mixing the base and curing agent at 10:1 ratio (SYLGARD 184 elastomer kit, Dow Corning). Bubbles were removed by centrifuging the PDMS mix for 2 min at 1000 rpm (5804 R, Eppendorf). After curing at room temperature for 4h, PDMS was spin coated for 30 s at 900 rpm on a patterned silicon wafer mould (microchannels of 80-120mm width and 50mm depth, described in Duda et al^10^) with spin coater (SPS Spin 150 and SPIN 150i). The PDMS-covered mould was then fully cured on a hotplate at 100°C for 10 min. The cured PDMS was then peeled off the wafer and stored between two transparency films (Nobo).

### Annular ablation of the columnar tissue

Annular ablations were conducted on a Zeiss LSM 880 confocal microscope with a pulsed Chameleon Vision II Ti:Sapphire laser (Coherent) tuned to 760 nm.

The protocol for apical annular ablations was as follows: A 512 × 512 pixel region in the middle of the wing disc pouch was imaged with a LD C-Apochromat 40x/1.1 water objective at 4x zoom. This region was chosen because it offered the flattest region of the wing disc and enabled maximising the size of the annular region to be ablated, i.e. inner and outer diameters of 186 and 249 pixels (19.30 and 25.84 μm). Each timepoint consisted of 3 z-planes 1 μm apart imaged at the fastest rate without time intervals, with the middle plane being the plane of ablation at the level of adherens junctions. 5 timepoints were captured prior to ablation and the tissue was imaged for a further 40 timepoints following ablation. The above settings allowed us to image the tissue every 1.66 ± 0.07 s, for a total duration of 75 s.

For analysis, 4D images of ablations (3D+time) were first deconvolved using Huygens Essential 23.10 (Scientific Volume Imaging). The EpiTools^49^ Matlab-based analysis module “Adaption Projection” was used with default parameters to obtain a single-plane timelapse image (Matlab R2014a, MathWorks). A custom-written code in Matlab then converted the Epitools output.vtk files to .tiffs. These images were then corrected for temporal drift/movement using “Linear Stack Alignment with SIFT”^50^ plugin mode in Fiji^51^. The outer boundary of the annular cut was fitted with an ellipse, and the aspect ratios extracted. For comparison between different conditions, we took the aspect ratio 60 s after ablation (first timepoint after 60 s) when the recoil plateaued.

### Annular ablation of the BM

Annular ablation of the BM was similar to those of the columnar tissue with these differences: The airyscan mode was used to acquire images. Each timepoint consisted of only 1 z-plane, allowing to image the tissue every 0.26 s. 5 timepoints were captured prior to ablation and the tissue was imaged for a further 150 timepoints following ablation, totalling to 39 s. To assess that the BM was cut successfully and not just bleached, a z-stack was taken imaging the BM (tagged with *vkg*-GFP) and the cells (tagged with *myr*-mRFP). In successful cuts, we observed that the cells whose BM was ablated extrude out of the BM (Fig. S2a). Therefore, we only included cuts where we could observe protruding cells on the basal side in the analysis.

For analysis, 3D images of ablations (2D+time) were first processed using Airyscan processing on Zen black (Zeiss) and then corrected for temporal drift/movement using “Linear Stack Alignment with SIFT”^50^ or “Template Matching and Slice Alignment”^52^ plugin mode in Fiji^51^. The outer boundary of the annular cut was fitted with an ellipse, and the aspect ratios extracted. For comparison between different conditions, we took the aspect ratio 37 s after ablation (first timepoint after 37 s) when the recoil plateaued.

Confirming that no new collagen IV is incorporated into the network in *ex vivo Drosophila* wing discs Experiments were conducted on a Nikon CSU-W1 SoRa microscope with a CFI Apochromat Lambda S 40XC water objective at 1x zoom. A 92 × 92 pixel (10 μm × 10 μm) region was bleached using the 488 laser. The wing disc was subsequently imaged every 30-60 min for up to 4 h.

For analysis, two 50 × 50 pixel (5.4 μm × 5.4 μm) regions were chosen outside and inside the bleached square (labelled as ctrl and bleached regions in Fig S2c) and the mean fluorescent intensity (*I_ctrl_* and *I_bleach_*) was calculated Fiji^51^. To quantify the background fluorescence intensity (*I_background_*), another region with the same size was chosen outside the wing disc. The normalised mean intensity (*I_normalised_*) of the ctrl and bleached regions were then calculated as below:

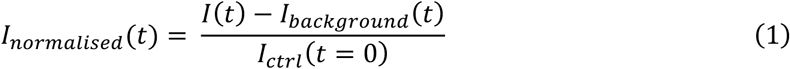

where *I* can be either *I_ctrl_* or *I_bleach_*.

### Quantifying the structure of collagen IV network

Experiments were conducted on a Nikon CSU-W1 SoRa microscope with a CFI Apochromat Lambda S 40XC water objective at 1x zoom. After acquiring an image at the anchor state, the wing disc was stretched and then imaged every hour for up to 4 h. Control experiments were conducted by imaging the wing disc anchored in the stretcher for 4 h.

The alignment of the collagen IV network was assessed using AFT (Alignment by Fourier Transform) workflow (the Matlab implementation)^36^. Images were first maximum projected in Fiji^51^ to obtain a single-plane timelapse image and corrected for temporal drift/movement using the “Template Matching and Slice Alignment”^52^ plugin. Next, a 250 × 250 pixel (27.08 μm × 27.08 μm) region was chosen in the middle of the wing disc pouch to conduct the AFT analysis. The AFT workflow includes some pre-built parameter search functions that allow for determining the length scale at which the difference between two conditions is greatest. We expected the difference between wing discs in anchor and immediately after (0 h) stretch to be the greatest. Therefore, we chose a test set of images from these two conditions and run the pre-built parameter search functions to determine the parameter set that gave the maximum difference between these conditions. This parameter set was then used for the final analysis: 29 pixel windows with a 50% overlap and neighbourhood radius of 6 pixels. The original code reports angles in the range [0 180), with 0° and 90° corresponding to alignment in the direction of the y and x axis respectively. A 90° rotation was therefore applied such that 0° and 90° correspond to alignment along the x and y axis respectively. The code was also modified to output the order maps.

For the purpose of visualisation of collagen IV alignment (and not the AFT analysis – Fig 2f,g, Fig S2d, Fig 4e,f), the images were first denoised using “denoise.ai” plugin in NIS Elements (Nikon) before maximum projecting and correcting for temporal drift/movement using “Template Matching and Slice Alignment”^52^ in Fiji^51^. Images where then cropped to a 250 × 250 pixel region, corresponding to the same region that was used in the alignment analysis above. Next, brightness was auto-adjusted for individual frames and applied to them individually.

### Statistical analysis

All curve fitting and plotting were conducted using custom-written code in MATLAB 2022.a (MathWorks) and Python (3.12.5). For each dataset, outliers were defined as the values that fell outside the range [q1 − w(q3 − q1), q3 + w(q3 − q1)], where q1 and q3 were the 25^th^ and 75^th^ percentiles of the data and w was 1.5. For all box plots, the edges of the box represent the 25^th^ and 75^th^ percentiles of the data, the red line shows the median and the whiskers extend to include the most extreme data points that are not considered to be outliers. Statistical analysis was performed in MATLAB, using paired two-sided Wilcoxon signed rank test where the same wing discs were analysed over time (Fig. 2h,i and 4g,h) or a two-sided Wilcoxon rank sum test elsewhere. Statistical differences are shown as follows: P < 0.01 (**) and 0.01< P < 0.05 (*) and represent the level of significance where we have a minimum of statistical power of 80%. Changes with P > 0.05 or where statistical power was less than 80% were considered non-significant. Actual P-values and statistical power are written in figure legends. The statistical power was calculated using G* Power^53^. Points/n-numbers on each box plot represent individual wing discs/simulations.

### Coarse-grained model of collagen IV network

#### Overview

We used a coarse-grained model for collagen IV, where each end of a collagen IV molecule is represented by a reactive bead which can bind to other molecule ends, and these ends are connected by a rod. The model is implemented in molecular dynamics using the software LAMMPS^54^. Full details of the model can be found in ref.^37^.

#### Simulation setup

A number of collagen IV molecules, *N*_7_, were initiated in a simulation box which was quasi-2D. The simulation length scale was σ and one molecule length was 3.5σ representing the end-to-end distance of a real molecule, ≈ 270 nm^37,55^. The simulations were performed in the canonical ensemble: a Velocity-Verlet integrator which was coupled to a Langevin thermostat at temperature *T* with a viscous friction coefficient γ = 0.1*m*/τ, where 2*m* is the mass of a molecule and 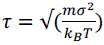 is the simulation time unit. The simulation timestep was Δt = 0.005 τ. In Fig. 3, 4a-d and S3d-e,g-j, *N_p_* = 15680 molecules were simulated in a box with dimensions *L_x_* = *L_y_* = 164σ, *L_z_* = 12σ. In Fig. S3a-c, to match the large length scale of experimental ablations, *N_p_* = 132845 molecules were simulated in a box with dimensions *L_x_*= *L_y_*= 470σ, *L_z_* = 12σ. The density of simulations is constant at 0.05 *mols*/σ^3^. The relevant parameters optimised and varied for this work are i) the bond length above which a bond between two protomers is considered to break, *D_break_* ii) the bond length below which a bond between to protomers may be made, *D_make_* iii) the probability to break, *P_break_*. and iv) make bonds, *D_make_*. These parameters control the amount of bond breaking and making occurring in a simulation and are shown in Table 3 below.

**Table 3.**
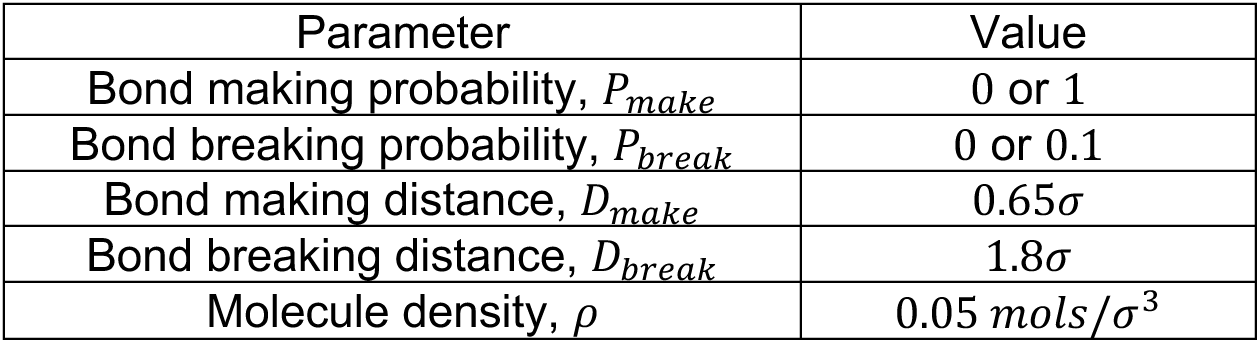
List of simulation parameters optimised for this work.

## Simulation protocols

In all simulations, protomers were first assembled into percolating networks over a time 2 × 10^6^ Δ*t* with parameters as in Table 3, *P_break_* = 0.1 and *P_make_* = 1. In Fig. 3 and 4, a uniaxial strain of 100% was applied to the networks in the x-direction over 0.2 × 10^6^Δ*t* and the networks subsequently equilibrated at this new strain which was kept fixed over 35 × 10^6^Δ*t*. In the ‘no remodelling’ condition, no bonds were made or broken in the time (*P_break_* = *P_make_* = 0). In the ‘remodelling’ condition, bonds were made with a probability *P_make_* = 1 and broken with a probability *P_break_* = 0.1. In Fig. S3c, the range of strains between 50%-150% were applied over a time 0.1 × 10^6^Δ*t*. In Fig. S3a the network relaxes at 100% stretch with no bond rearrangement. To simulate annular ablations in Fig. S3b-c, a number of molecules and their bonds were removed from the simulation in the shape of an annulus, with inner radius of 69σ and outer radius of 93σ and the resulting perturbed network rearranged in response.

### Calculating proportion of fluorescently-tagged to non-tagged molecules in simulations

*Drosophila* has two collagen IV genes, Col4α1 (*Cg25c*) and Col4α2 (*vkg*), which encode for collagen IV α1 and α2 chains respectively. Each collagen IV molecule is formed by the assembly of two α1 and one α2 chains. In the experiments looking at the nematic order of collagen IV network, we used overexpression lines of Col4α1-mScarlet. This means that only part of the collagen IV network is visualised. To incorporate a similar condition in our simulations, we estimated the proportion of fluorescently-tagged collagen IV molecules in our experiments as below:

Assume we have *m* number of wild-type collagen IV molecules. Since each collagen IV molecule has 2 two α1 chains, this gives us 2*m* endogenous α1 chains. We further assume that we have *n* number of overexpressed fluorescently-tagged α1 chains. The total number of α1 chains in our system would therefore be: 2*m* + *n*. To form a collagen IV molecule, we need to choose 2 chains from the mixed pool of wild-type and fluorescently-tagged α1 chains. Therefore, the total number of combinations we can have is:

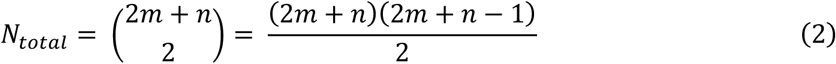

This includes molecules that have either none, one or two fluorescently-tagged α1 chains. Having one fluorescently-tagged α1 chains will make the molecule fluorescent (and therefore visualised). To calculate the probability of having a fluorescently-tagged molecule, we do as follows:

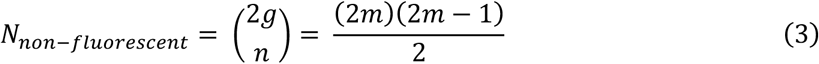

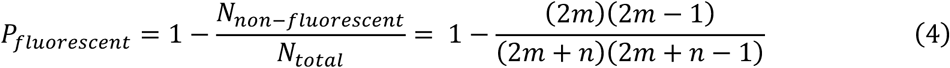

Assuming that we m and n are infinite:

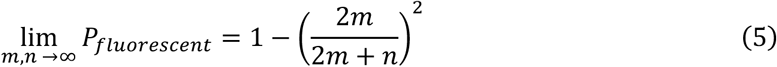

Since experimentally we don’t know the number of over-expressed Col4α1-mScarlet genes, we consider two scenarios:

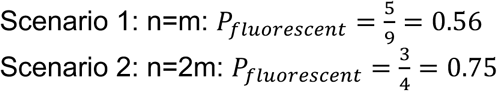

In the simulations, this means that if we have 100 non-tagged molecules, we should have:

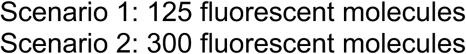

In the case of 7S mutations, these fluorescent molecules will also be mutant.

### Matching the extent of deformation in simulations with experiments

To match the extent of deformation and timescales of simulations with experiments, we compared the response of a stretched BM to annular ablations in the two settings. Following ablation of a stretched BM network, the cut region recoils more in the direction of stretch, leading to an increase in the aspect ratio of the cut boundary. As in the experiments, we quantify the aspect ratio of the outer boundary of the cut by fitting an ellipse to it. The aspect ratio increases over time and eventually plateaus at a timescale of τ_2_∼ 1.5 × 10^6^ Δ*t* after ablation, which corresponds to the minute-timescale plateau observed experimentally (Fig S2b). Since the nematic order changes in the timescale of hours, we ensured these simulations are run for timescales larger than that of annular ablations (i.e. τ_2_).

To match the extent of deformation in simulations with experiments, we performed *in silico* annular ablations applying different strains and compared their final aspect ratio with experiments. We chose 100% as our applied strain for all subsequent simulations, as its aspect ratio fell within our experimental range (Fig S3c).

### Post-processing of simulation snapshots for analysis

To compare simulation snapshots with microscopy images, the following post-processing was performed on the 7S beads that are visualised in the snapshots: A Gaussian point-spread function was applied to the 7S ends of a fraction of the molecules. The width of the point spread function was 3.5σ, representing the 200-250 nm resolution limit of confocal microscopy. The images were then visualised as a heatmap of these summed point-spread functions. In Fig 3 and S3d-e, the maximum brightness was capped at 30 for unstretched networks and 20 for stretched networks, such that the more diluted stretched networks could be visualised in detail. In Fig. 4a-c, the max brightness was capped at 30 for unstretched networks, 17 for stretched networks where 75% of protomers were mutant and 13 for stretched networks where 56% of protomers were mutant.

The AFT workflow (the Matlab implementation)^36^ was used with the following parameters: 54 pixel windows with a 50% overlap and neighbourhood radius of 4 pixels. The data was first smoothed using moving average with a sliding window size of 0.404 × 10^6^ Δ*t* and then binned into relevant intervals. The data points in each bin were then averaged to get a single nematic order per bin per wing disc. The stress is a direct readout from LAMMPS and the alignment was calculated as shown in Fig. 3j.

## Figures

**Fig S1:**
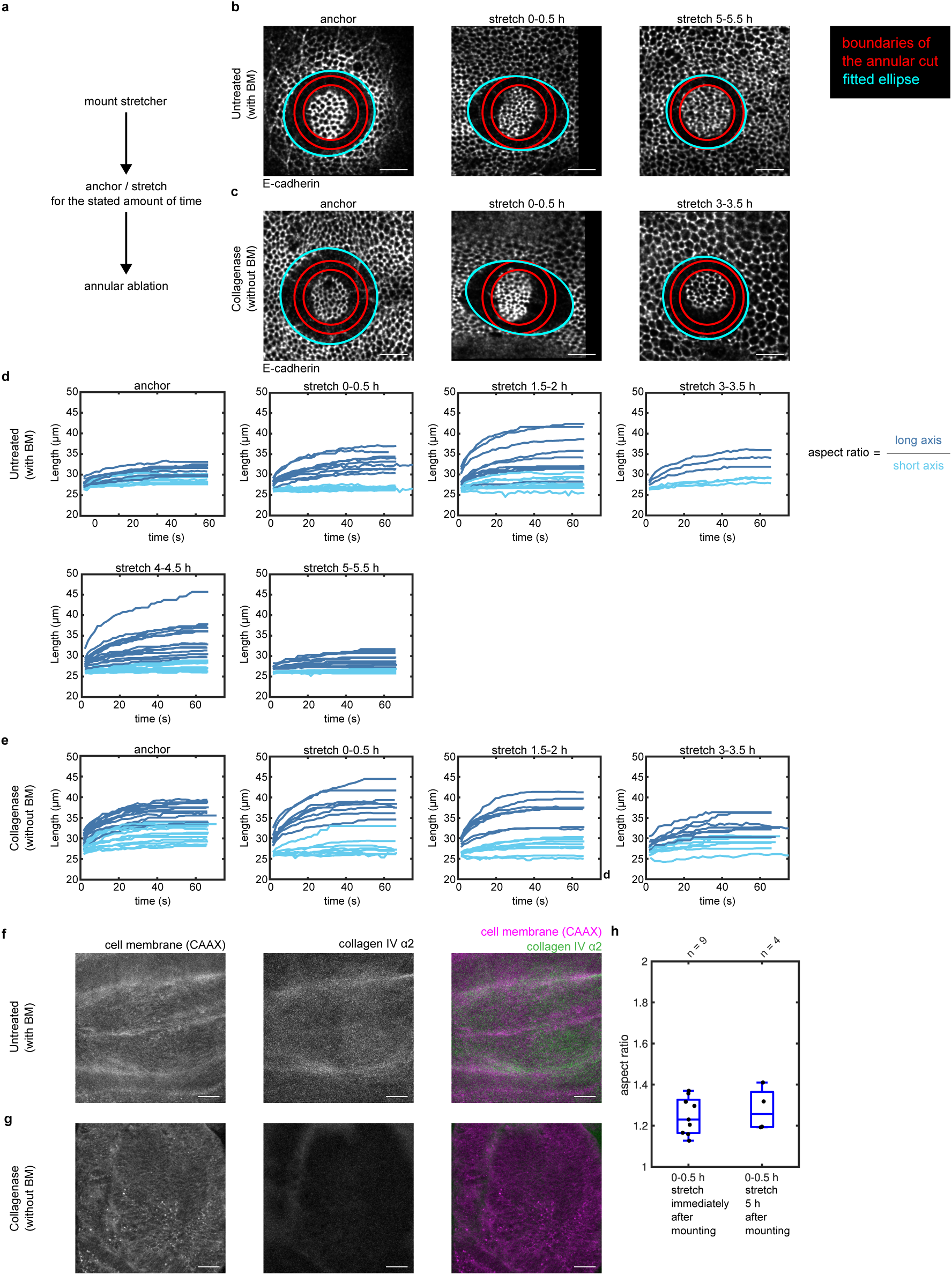
Details of the experimental protocol and quantification of annular ablation of the tissue. (a) Schematic diagram detailing the steps of the annular ablation experiments. (b,c) Snapshots of the annular ablation experiments shown in Fig 1e,h overlayed with fitted ellipses used for aspect ratio quantifications. The area between the two red circles shows the ablated region. The fitted ellipse to the outer boundary of the cut is shown in cyan. Scale bars = 10 μm. (d,e) Temporal change in the long (dark blue) and short (light blue) axis of the fitted ellipses for different anchor/stretch conditions in untreated (d) and collagenase-treated(e) wing discs. (f,g) Control experiments to test the effectiveness of the collagenase treatment. The collagen IV (Col4α2) signal almost disappears after collagenase-treatment (compare middle panels of f and g), whereas the membrane marker (CAAX) remains unchanged (left panels of f and g). Scale bars = 20 μm. (h) Control experiment to test viability of wing discs in the stretcher up to 5 h (n.s., P = 0.6).

**Fig S2:**
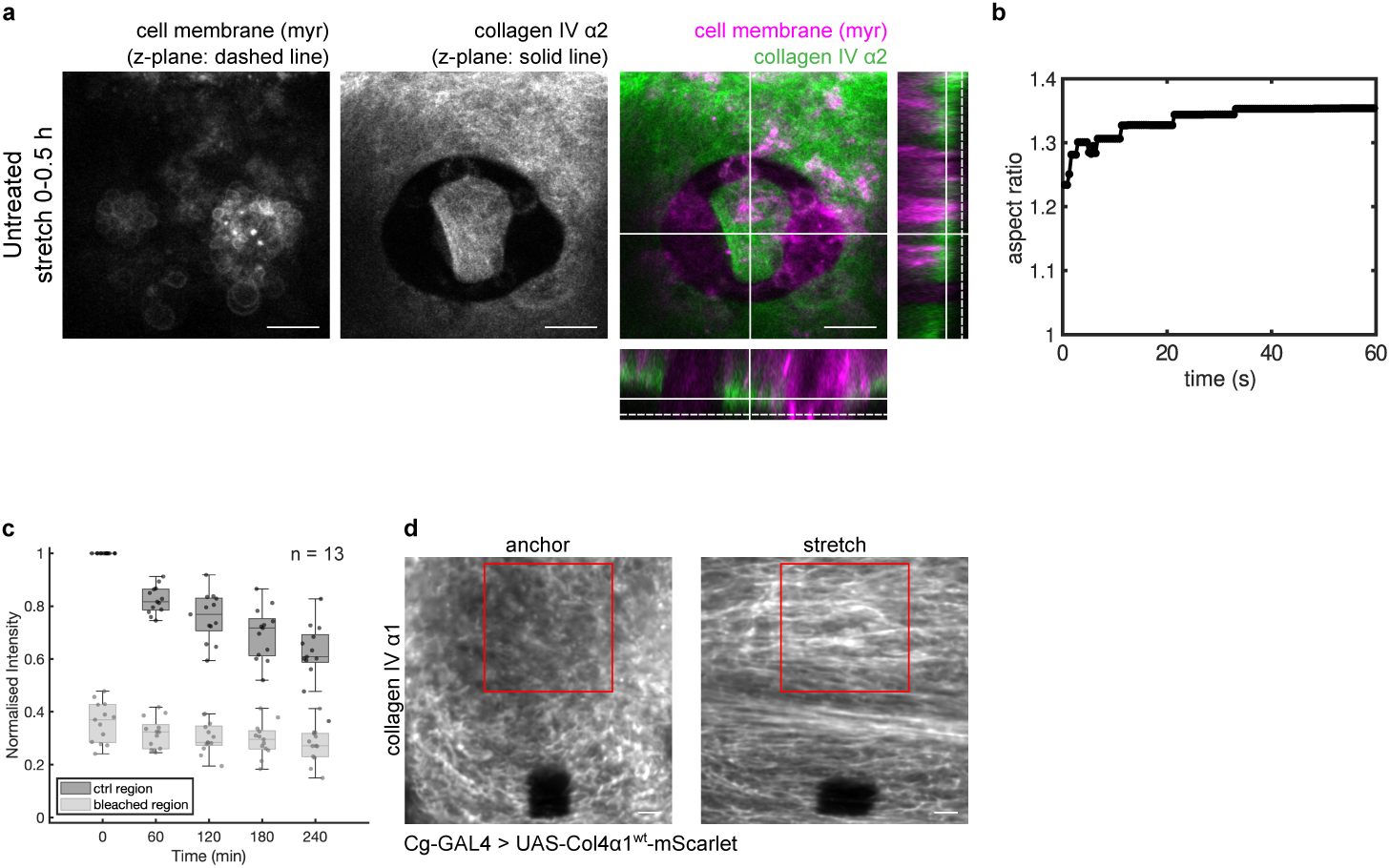
Details of annular ablation experiments and imaging of the BM. (a) Snapshot of an annular ablation experiment as shown in Fig 2b,c. The panels show the cell membranes (*myr*, left), BM (Col4α2, middle) and their overlay (right). The cells extruded from the annular cut after ablation are shown in the left panel and their corresponding z-plane is shown as a dashed line in the right panel. The collagen IV network is shown in the middle panel and its corresponding z-plane is shown as a solid line in the right panel. Scale bars = 10 μm. (b) Example of the temporal evolution of aspect ratio of a BM annular cut. (c) Control experiment to show there is no addition of new material in the BM of dissected wing discs. (d) Zoomed-out near-super-resolution confocal images of a wing disc before (left) and after (right) stretch. The ROI used for nematic order calculations and shown in Fig 2g is highlighted in red. Scale bars = 5 μm.

**Fig S3:**
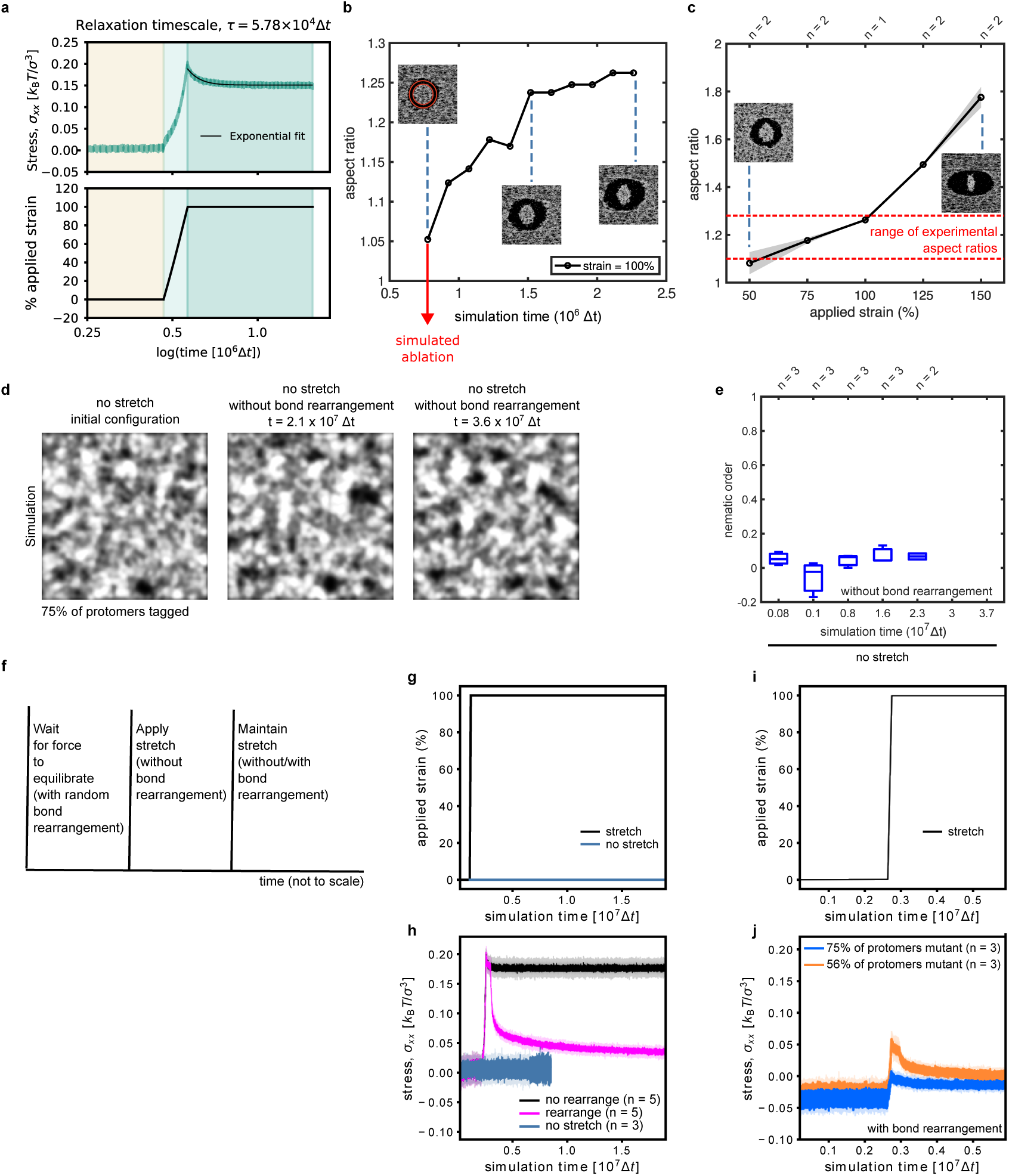
Details of simulated collagen IV stretching experiments with additional quantifications. (a) The temporal evolution of stress before (cream), immediately (light green) and after (dark green) application of 100% strain without bond rearrangement. There is a fast relaxation immediately after stretch that can be fitted with an exponential function to calculate its characteristic time. (b) Temporal evolution of outer boundary of a simulated annular cut immediately after application of 100% strain. The insets show snapshots at different timepoints. The area between the two red circles in the left inset shows the initial cut region. (c) The change in final aspect ratio of the annular cut in response to different amounts of applied strain. The region between the two red dashed lines shows the range of experimental aspect ratios. (d) Snapshot of a simulations with normal protomers and without stretch over time. Only 75% of the protomers are tagged to replicate experimental conditions. (e) Box plot showing temporal evolution of the nematic order of simulated networks in the absence of external deformation (i.e. no stretch). (f) The protocol outlining different steps of a stretching simulation. (g,h) Temporal evolution of the applied strain (g) and its respective stress for simulations with normal protomers. The solid lines in (h) show average of several simulations and the shadowed regions show their standard deviation. (i,j) Temporal evolution of the applied strain (i) and its respective stress for simulations with mutated protomers. Two conditions were tested with different concentration of mutated protomers. The solid lines in (j) show average of several simulations and the shadowed regions show their standard deviation.

## Data and code availability

Data and code will be available upon request.

## Acknowledgements

We thank R. Barrientos for critical reading of this work, and B. Sánchez Sánchez and B. Stramer for kindly sharing fly lines. We thank the National Centre for the Replacement, Refinement and Reduction of Animals in Research (NC3Rs) for funding this work (NC/T002425/1 and an Early Career Engagement Award to N.K.). N.K. was also supported by a Leverhulme Trust project grant (RPG-2020-068) and an MRC Fellowship (MR/W027437/1) awarded to Y.M., and a Leverhulme Trust project grant (RPG-2024-147). Y.M. was funded by MRC award MR/W027437/1, a Lister Institute Research Prize and EMBO Young Investigator Programme. This work has also received funding from the European Research Council (ERC) under the European Union’s Horizon 2020 research and innovation programme (grant agreement No. 802960; B.M., A.Š.). A.Š was also funded by the EMBO Young Investigator Programme and the Vallee Foundation.

## Author contributions

N.K. and Y.M. conceived the project. N.K. designed, performed and analysed experiments. B.M. and A.Š. designed and developed the computational model, with feedback from N.K. and Y.M. N.K. B.M. and Y.M. wrote the manuscript with contributions from A.Š. All authors discussed the results and the manuscript.

## References

1. He, Z., Ritchie, J., Grashow, J. S., Sacks, M. S. & Yoganathan, A. P. In Vitro Dynamic Strain Behavior of the Mitral Valve Posterior Leaflet. Journal of Biomechanical Engineering 127, 504–511 (2005).

2. Perlman, C. E. & Bhattacharya, J. Alveolar expansion imaged by optical sectioning microscopy. Journal of Applied Physiology 103, 1037–1044 (2007).

3. Maiti, R. et al. In vivo measurement of skin surface strain and sub-surface layer deformation induced by natural tissue stretching. Journal of the Mechanical Behavior of Biomedical Materials 62, 556–569 (2016).

4. Bhattacharya, D., Zhong, J., Tavakoli, S., Kabla, A. & Matsudaira, P. Strain maps characterize the symmetry of convergence and extension patterns during zebrafish gastrulation. Sci Rep 11, 19357 (2021).

5. Blanchard, G. B. et al. Tissue tectonics: morphogenetic strain rates, cell shape change and intercalation. Nature methods 6, 458–64 (2009).

6. Sacks, M. S. et al. In-Vivo Dynamic Deformation of the Mitral Valve Anterior Leaflet. The Annals of Thoracic Surgery 82, 1369–1377 (2006).

7. Wyatt, T., Baum, B. & Charras, G. A question of time: tissue adaptation to mechanical forces. Current Opinion in Cell Biology 38, 68–73 (2016).

8. Harris, A. R. et al. Characterizing the mechanics of cultured cell monolayers. Proceedings of the National Academy of Sciences of the United States of America 109, 16449–16454 (2012).

9. Khalilgharibi, N. et al. Stress relaxation in epithelial monolayers is controlled by the actomyosin cortex. Nature Physics 15, 839–847 (2019).

10. Duda, M. et al. Polarization of Myosin II Refines Tissue Material Properties to Buffer Mechanical Stress. Developmental Cell 48, 245–260.e7 (2019).

11. Etournay, R. et al. Interplay of cell dynamics and epithelial tension during morphogenesis of the Drosophila pupal wing. eLife 4, e07090 (2015).

12. Collinet, C., Rauzi, M., Lenne, P. & Lecuit, T. Local and tissue-scale forces drive oriented junction growth during tissue extension. Nature Cell Biology 17, 1247–1258 (2015).

13. Lye, C. M. et al. Mechanical Coupling between Endoderm Invagination and Axis Extension in Drosophila. PLoS Biol 13, e1002292 (2015).

14. Campinho, P. et al. Tension-oriented cell divisions limit anisotropic tissue tension in epithelial spreading during zebrafish epiboly. Nature Cell Biology 15, 1405–1414 (2013).

15. Wyatt, T. P. J. et al. Emergence of homeostatic epithelial packing and stress dissipation through divisions oriented along the long cell axis. Proceedings of the National Academy of Sciences 112, 5726–5731 (2015).

16. Khalilgharibi, N. & Mao, Y. To form and function: on the role of basement membrane mechanics in tissue development, homeostasis and disease. Open Biol. 11, 200360 (2021).

17. Page-McCaw, A. & Ferrell, N. Basement membrane structure and function: Relating biology to mechanics. Matrix Biology 141, 16–31 (2025).

18. Chaudhuri, O., Cooper-White, J., Janmey, P. A., Mooney, D. J. & Shenoy, V. B. Effects of extracellular matrix viscoelasticity on cellular behaviour. Nature 584, 535–546 (2020).

19. Yurchenco, P. D. & Schittny, J. C. Molecular architecture of basement membranes. The FASEB Journal 4, 1577–1590 (1990).

20. Chlasta, J. et al. Variations in basement membrane mechanics are linked to epithelial morphogenesis. Development 144, 4350–4362 (2017).

21. Crest, J., Diz-Muñoz, A., Chen, D.-Y., Fletcher, D. A. & Bilder, D. Organ sculpting by patterned extracellular matrix stiffness. eLife 6, 1–16 (2017).

22. Harmansa, S., Erlich, A., Eloy, C., Zurlo, G. & Lecuit, T. Growth anisotropy of the extracellular matrix shapes a developing organ. Nat Commun 14, 1220 (2023).

23. Guerra Santillán, K. Y., Dahmann, C. & Fischer-Friedrich, E. Elastic Contractile Stress in the Basement Membrane Generates Basal Tension in Epithelia. PRX Life 2, 013004 (2024).

24. Fabris, G. et al. Nanoscale Topography and Poroelastic Properties of Model Tissue Breast Gland Basement Membranes. Biophysical Journal 115, 1770–1782 (2018).

25. Nam, S., Lee, J., Brownfield, D. G. & Chaudhuri, O. Viscoplasticity Enables Mechanical Remodeling of Matrix by Cells. Biophysical Journal 111, 2296–2308 (2016).

26. Wisdom, K. M. et al. Matrix mechanical plasticity regulates cancer cell migration through confining microenvironments. Nature Communications 9, (2018).

27. Forgacs, G., Foty, R. A., Shafrir, Y. & Steinberg, M. S. Viscoelastic Properties of Living Embryonic Tissues: a Quantitative Study. Biophysical Journal 74, 2227–2234 (1998).

28. Doubrovinski, K., Swan, M., Polyakov, O. & Wieschaus, E. F. Measurement of cortical elasticity in *Drosophila melanogaster* embryos using ferrofluids. Proceedings of the National Academy of Sciences 114, 1051–1056 (2017).

29. Khalilgharibi, N. et al. Stress relaxation in epithelial monolayers is controlled by the actomyosin cortex. Nat. Phys. 15, 839–847 (2019).

30. Bonnet, I. et al. Mechanical state, material properties and continuous description of an epithelial tissue. Journal of The Royal Society Interface 9, 2614–2623 (2012).

31. Leslie, H., Sperling, S. Introduction to Physical Polymer Science. (Wiley-Blackwell, 2015).

32. McCall, P. M., MacKintosh, F. C., Kovar, D. R. & Gardel, M. L. Cofilin drives rapid turnover and fluidization of entangled F-actin. Proc. Natl. Acad. Sci. U.S.A. 116, 12629–12637 (2019).

33. Iyer, K. V., Piscitello-Gómez, R., Paijmans, J., Jülicher, F. & Eaton, S. Epithelial Viscoelasticity Is Regulated by Mechanosensitive E-cadherin Turnover. Current Biology 29, 578–591.e5 (2019).

34. Barrientos R et al. Basement membrane turnover controls cell shape.

35. Pastor-Pareja, J. C. & Xu, T. Shaping Cells and Organs in Drosophila by Opposing Roles of Fat Body-Secreted Collagen IV and Perlecan. Developmental Cell 21, 245–256 (2011).

36. Marcotti, S. et al. A Workflow for Rapid Unbiased Quantification of Fibrillar Feature Alignment in Biological Images. Frontiers in Computer Science 3, 1–16 (2021).

37. Meadowcroft, B. et al. Nonequilibrium remodeling of collagen IV networks in silico. PRX Life (2025) doi:10.1103/gdd5-rnh7.

38. Serna-Morales, E. et al. Extracellular matrix assembly stress initiates Drosophila central nervous system morphogenesis. Developmental Cell 58, 825–835.e6 (2023).

39. Tozluoǧlu, M., et al. Planar Differential Growth Rates Initiate Precise Fold Positions in Complex Epithelia. Developmental Cell 51, 299–312.e4 (2019).

40. Khalilgharibi, N., Fouchard, J., Recho, P., Charras, G. & Kabla, A. The dynamic mechanical properties of cellularised aggregates. Current Opinion in Cell Biology 42, 113–120 (2016).

41. Chang, J. & Chaudhuri, O. Beyond proteases: Basement membrane mechanics and cancer invasion. Journal of Cell Biology 218, 2456–2469 (2019).

42. Wershof, E. et al. A FIJI macro for quantifying pattern in extracellular matrix. Life Science Alliance 4, e202000880 (2021).

43. Harunaga, J. S., Doyle, A. D. & Yamada, K. M. Local and global dynamics of the basement membrane during branching morphogenesis require protease activity and actomyosin contractility. Developmental Biology 394, 197–205 (2014).

44. Soh, A. W. J. et al. On-demand delivery of fibulin-1 protects the basement membrane during cyclic stretching in C. elegans. Developmental Cell (2025) doi:10.1016/j.devcel.2025.07.015.

45. Peebles, K. E. et al. Peroxidasin is required for full viability in development and for maintenance of tissue mechanics in adults. Matrix Biology 125, 1–11 (2024).

46. Keeley, D. P. et al. Comprehensive Endogenous Tagging of Basement Membrane Components Reveals Dynamic Movement within the Matrix Scaffolding. Developmental Cell 17, 320–329 (2020).

47. Huang, J., Zhou, W., Dong, W., Watson, A. M. & Hong, Y. Directed, efficient, and versatile modifications of the *Drosophila* genome by genomic engineering. Proc. Natl. Acad. Sci. U.S.A. 106, 8284–8289 (2009).

48. Martin, A. C., Kaschube, M. & Wieschaus, E. F. Pulsed contractions of an actin–myosin network drive apical constriction. Nature 457, 495–499 (2009).

49. Heller, D. et al. EpiTools: An Open-Source Image Analysis Toolkit for Quantifying Epithelial Growth Dynamics. Developmental Cell 36, 103–116 (2016).

50. Lowe, D. G. Distinctive Image Features from Scale-Invariant Keypoints. International Journal of Computer Vision 60, 91–110 (2004).

51. Schindelin, J., et al. Fiji: an open-source platform for biological-image analysis. Nature Methods 9, 676–682 (2012).

52. Tseng, Q., et al. Spatial Organization of the Extracellular Matrix Regulates Cell-Cell Junction Positioning. Proceedings of the National Academy of Sciences of the United States of America vol. 109 (2012).

53. Faul, F., Erdfelder, E., Lang, A.-G. & Buchner, A. G*Power 3: A flexible statistical power analysis program for the social, behavioral, and biomedical sciences. Behavior Research Methods 39, 175–191 (2007).

54. Thompson, A. P. et al. LAMMPS - a flexible simulation tool for particle-based materials modeling at the atomic, meso, and continuum scales. Computer Physics Communications 271, 108171 (2022).

55. Al-Shaer, A. et al. Sequence-dependent mechanics of collagen reflect its structural and functional organization. Biophysical Journal 120, 4013–4028 (2021).

